# Oxidized products of *α*-linolenic acid negatively regulate cellular survival and motility of breast cancer cells

**DOI:** 10.1101/517094

**Authors:** Jorge L. Gutierrez-Pajares, Celine Ben Hassen, Camille Oger, Jean-Marie Galano, Thierry Durand, Philippe G. Frank

**Author notes:** **Address for reprint requests and other correspondence:** J.L. Gutierrez-Pajares or P.G. Frank, Université de Tours, INSERM, UMR1069, Nutrition, Croissance et Cancer, Tours, France. or.

## Abstract

**Background:** Cancer is a major cause of death in the world, and more than six million new cases are reported every year. Despite recent advances in our understanding of the biological processes leading to the development and progression of cancer, there is still a need for new and effective agents to treat this disease. Phytoprostanes (PhytoPs) and phytofurans (PhytoFs) are non-enzymatically oxidized products of α-linolenic acid that are present in seeds and vegetable oils. They have been shown to possess anti-inflammatory and apoptosis-promoting activities in macrophages and leukemia cells, respectively.

**Methods:** In this work, seven PhytoPs (PP1-PP7) and one PhytoFs (PF1) were evaluated for their cytotoxic, chemosensitization and anti-migratory activities using the MCF-7 and MDA-MB-231 breast cancer cell lines.

**Results:** Among the compounds tested, only three PhytoPs had a significant effect on cell viability compared to the control group: *Ent*-9-L_1_-PhytoP (PP6) decreased cell viability in both cell lines, while 16-F_1t_-PhytoP (PP1) and 9-L_1_-PhytoP (PP5) decreased viability in MCF-7 and MDA-MB-231 cells, respectively. When combined with a sub-cytotoxic dose of doxorubicin, these three PhytoPs significantly enhanced the cytotoxic effect on MCF-7 cells while the chemotherapeutic drug alone had no effect. In cellular motility assays, *Ent*-9-(*RS*)-12-*epi*-ST-Δ^10^-13-PhytoF could significantly inhibit cellular migration of MDA-MB-231 cells in a wound-healing and a transwell assays. In addition, *Ent*-9-(*RS*)-12-*epi*-ST-Δ^10^-13-PhytoF also enhanced cellular adhesion of MDA-MB-231 cells.

**Conclusions:** This study shows for the first time that the plant-derived compounds PhytoPs and PhytoFs could be further exploited alone or in combination with chemotherapy to improve the arsenal of therapies available against breast cancer.

## Background

Cancer is a major cause of death in the world, and while 18.1 million new cases are expected to be detected in 2018, 9.6 million cancer-related deaths may occur [1]. Despite recent advances in our understanding of the biological processes leading to the development of cancer, there is still a need for new and effective methods of treatment for this disease. Natural products and their derivatives represent an important source of new therapeutic agents, as a tremendous chemical diversity is found in millions of species of plants, animals, and microorganisms. Plant-derived compounds have played an important role in the development of several clinically effective anti-cancer agents [2]. Although a number of anticancer agents derived from plants have been identified, many compounds that may exhibit anti-cancer properties remain to be identified.

Phytoprostanes (PhytoPs) are prostaglandin-like compounds that are found in seeds and vegetable oils [3, 4]. Arachidonate (ARA, C20:4 n-6) may be transformed via a non-enzymatic oxidative cyclization into isoprostanes (IsoPs) in animals. In plants, on the other hand, α-linolenic acid (ALA, C18:3 n-3) oxidative cyclization leads to a series of PhytoPs and phytofurans (PhytoFs) [5]. Under high levels of oxygen tension, PhytoFs are preferentially formed, as compared to PhytoPs [6].

Previous studies have shown that IsoPs may represent markers of oxidative stress but may also have important physiological functions in obesity, cardiovascular diseases, and cancer. Clinical studies have shown that urinary levels of F_2_-IsoPs are correlated with increased risk of developing several types of cancer [7]. In one study, it was shown that for a BMI<23, elevated F_2_-IsoPs levels were associated with reduced risk of breast cancer. On the other hand, in patients with BMI>25, elevated levels of F_2_-IsoPs were associated with increased risk of breast cancer. However, whether these products are formed as a consequence or as a cause of the disease has not yet been established [7].

Since PhytoPs and PhytoFs are structurally similar to IsoPs and prostanoids, their biological activities are currently under investigation. It has been suggested that PhytoPs can trigger a defensive mechanism in plants [8]. Therefore, in *Arabidopsis thaliana*, exposure of leaves to PhytoPs mediates oxidative stress to induce the expression of proteins involved in stress response and redox regulation [9]. In mammals, PhytoPs have been shown to possess anti-inflammatory and cell-death promoting activities. It has also been shown that PhytoPs regulate cytokine production of inflammatory cells [10, 11] and down-regulate NF-κB in murine macrophages [11]. Moreover, PhytoPs have been reported to induce apoptosis in the leukemia Jurkat cell line [12]. The differential effect of PhytoPs and PhytoFs has been suggested to depend on the stereochemistry of the studied compound [4]. Importantly, IsoPs have been demonstrated to inhibit VEGF-induced migration of endothelial cells [13]. Minghetti *et al.* reported that 16-B_1_-PhytoP and 9-L_1_-PhytoPs protect immature neuronal cells from oxidative stress and promote differentiation of oligodentrocytes [14]. In addition, a diet enriched in flaxseed oil is associated with increased plasma levels of PhytoPs in healthy volunteers [15]. Importantly, studies have shown that, in breast cancer, ALA limits tumor progression by inhibiting cancer cell proliferation [16, 17] and tumor progression [18, 19]. These effects may be mediated by other n-3 fatty acids (EPA and DHA) but independent roles for ALA have been proposed [20, 21]. Oxygenated metabolites of ALA such as PhytoPs and PhytoFs may play a role in the inhibition of breast cancer progression mediated by ALA. Taken together, these data reinforce the need to understand the effects of PhytoPs and PhytoFs in health and disease.

In the present work, seven PhytoPs (PP1-PP7) and one PhytoF (PF1) were tested on breast cancer cell lines to determine their effect on cell viability and motility.

## Material and Methods

### Materials

Cell culture reagents were obtained from Fischer Scientific (Illkirch-Graffenstaden, France). Doxorubicin was obtained from Sigma-Aldrich (Saint-Quentin Fallavier, France). All other reagents were analytical grade.

### Cell lines

MCF-7 and MDA-MB-231 cells were obtained from the American Tissue Culture Collection (ATCC) (Molsheim, France). MDA-MB-231 and MCF-7 cells were grown in Dulbecco’s modified Eagle’s media (DMEM) containing 10% Fetal Bovine Serum (FBS) and 1% penicillin and streptomycin (complete media) in a humidified incubator kept at 37ºC with 5% CO_2._

### Source of ALA derivatives

Synthesis of PhytoPs and PhytoF was performed as previously described [22-24]. In the present study the following compounds were tested: 16-F_1t_-PhytoP (**PP1**), 16-*epi*-16-F_1t_-PhytoP (**PP2**), 16-B_1_-PhytoP (**PP3**), *Ent*-16-B_1_-PhytoP (**PP4**), 9-L_1_-PhytoP (**PP5**), *Ent*-9-L_1_-PhytoP (**PP6**), 9-E_1_-PhytoP (**PP7**) and *Ent*-9-(*RS*)-12-*epi*-ST-Δ^10^-13-PhytoF (**PF1**). The structure of these compounds is presented in Figure 1.

**Figure 1:**
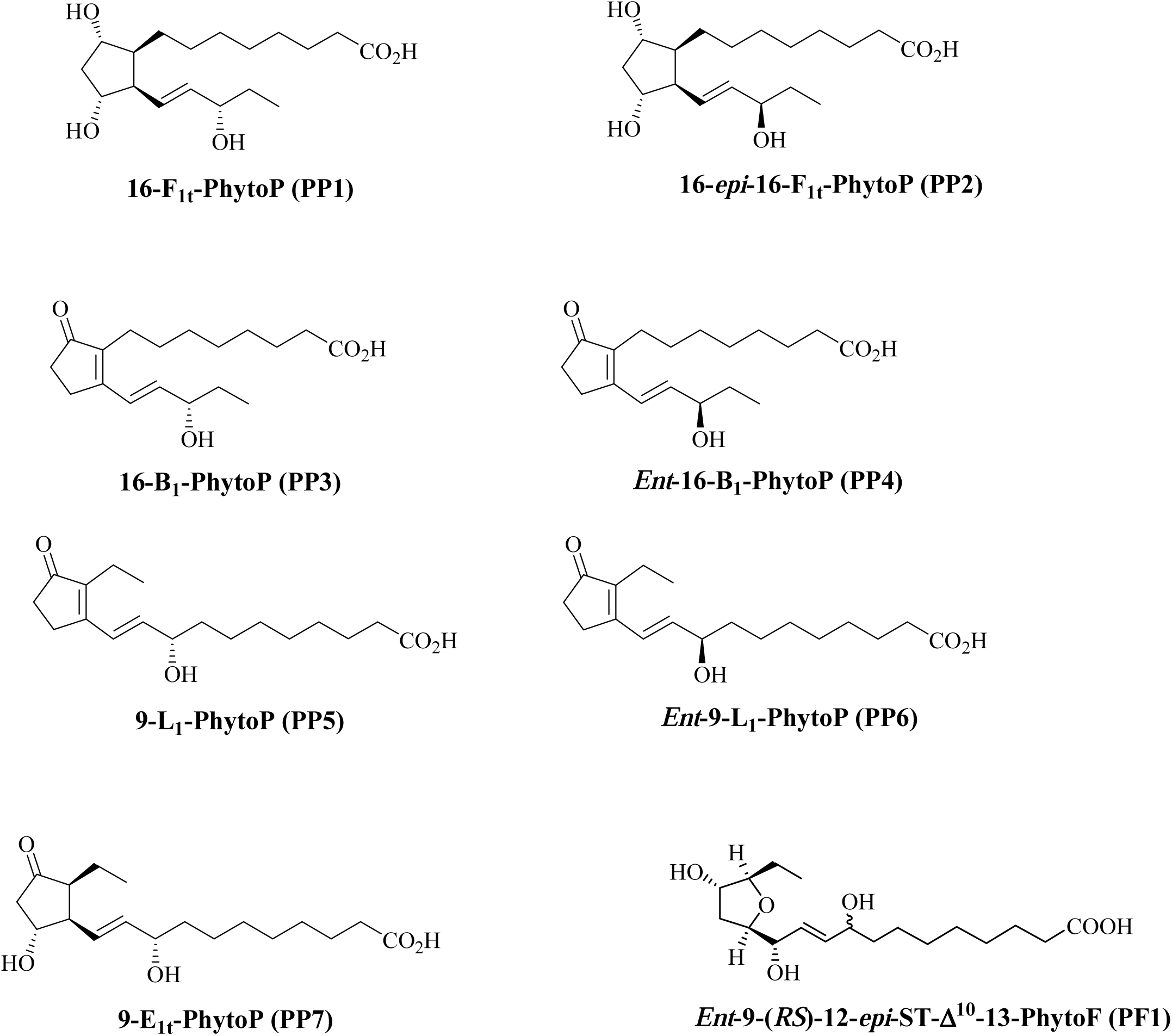
Structure of the phytoprostanes used in this study.

### Cellular density assay

Cells were seeded in 96-well plates in the media indicated for each experimental condition. At the end of the incubation with or without treatment, culture media was removed and a crystal violet (CV) staining solution was added to fix and stain cells for 1 h. After removal of the staining solution, plates were washed several times with distilled water to remove excess stain. Plates were subsequently dried at 37°C for at least 3 h. Cellular stain was recovered by adding a 10% acetic acid solution and quantified by spectrophotometry.

### Cell cycle analysis

For each study, cells were detached with trypsin, fixed in 70% ethanol, treated with RNAse (5 µg/mL), and stained with propidium iodide (3 µM). Analyses were performed with a Gallios flow cytometer (Beckman Coulter). Gating and percentage of cells for cell cycle phases were determined using the Kaluza® Analysis software v1.3 (Beckman Coulter).

### Cellular migration assay

MDA-MB-231 cells were seeded in a 96-well plate at 12.5×10^3^ cells/well in complete media. After the culture had reached confluency, a wound was applied with a 10 µl-pipette tip to each well. Cells were washed twice with PBS containing calcium and magnesium, and culture media (95 µL) with 0.1% fatty-acid-free BSA (faf-BSA) with or without added PhytoPs or PhytoF. FBS (5 µL) was directly added to each well 30 minutes later. Pictures (3 per well) with a 10x magnification were taken 24 h later. Cell-free area was determined for each picture and averaged per well.

### Cell adhesion

Cells were gently detached with Accutase® (Sigma-Aldrich), counted, resuspended in 0.1% fatty acid free-BSA media, and seeded (2×10^4^ cells/50µL/well) in 96-well plates that were pre-coated or not with extracellular matrix from complete media, blocked with 0.5% BSA for 1 h, and washed twice with PBS before seeding cells. Cells were incubated for 1 h at 37 °C and then washed and stained with a crystal violet solution, as described above.

### Statistical analysis

Data were expressed as mean ± standard error of the mean. Data were analyzed with Kruskal-Wallis nonparametric ANOVA followed by the Dunn’s multiple comparison test. Statistical significance was established at *p* < 0.05 level. Analyses were performed using GraphPad Prism v6.0.

## Results

### PP6 reduces cellular survival of both MCF-7 and MDA-MB-231 cells

The present study was undertaken to determine the role of several members of the PhytoP family (**Figure 1**) on breast tumor cell properties. In the first series of experiments, we analyzed their effects on cellular survival in the presence of 1% FBS for 48 h (**Figure 2A**). Our data show that PP1, PP6, and PP7 reduced survival of MCF-7 cells, while PP5 and PP6 had similar effects on MDA-MB-231 cells (Figure 2B). Interestingly, PP3 slightly promoted proliferation of MCF-7 but had limited effects on MDA-MB-231 cells. Therefore, PP6 was the only PhytoP to display cytotoxic activity against both cell lines.

**Figure 2:**
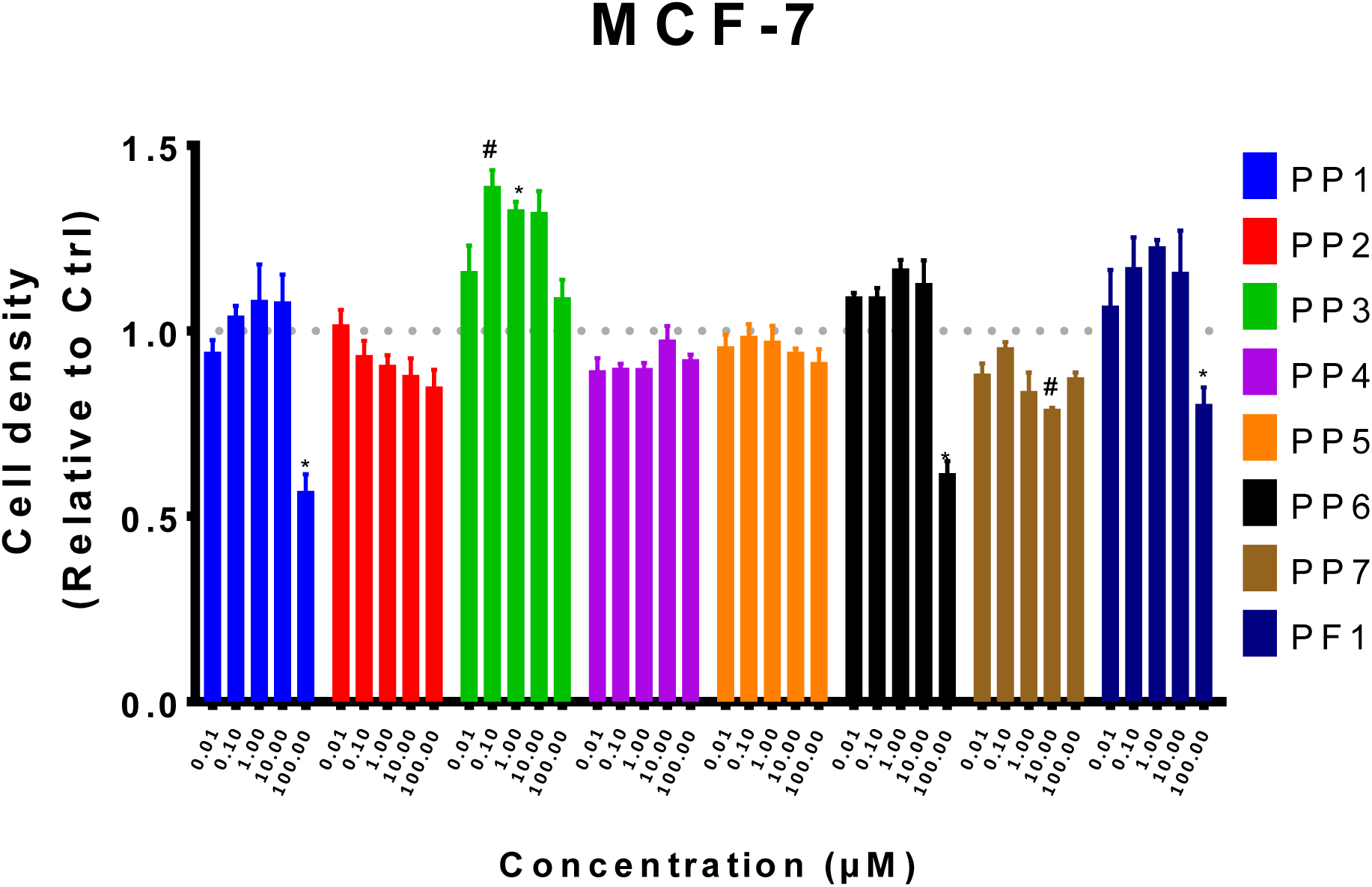
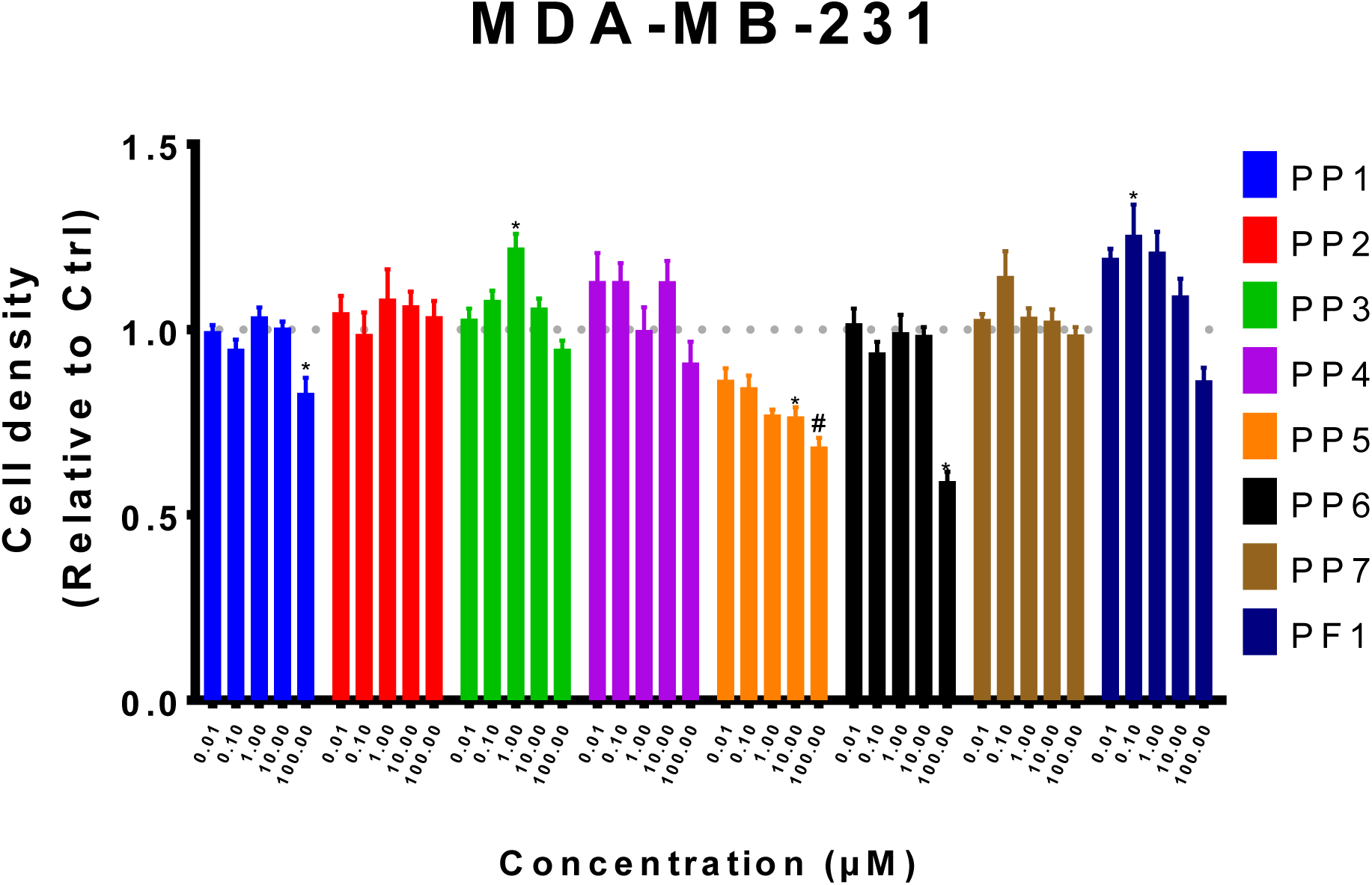
PP6 reduces cell survival of both MCF-7 and MDA-MB-231 cells. MCF-7 (**A**) and MDA-MB-231 (**B**) were seeded in 96-well plates and treated with increasing concentrations (0.01-100 μM) of PhytoPs (PP1-7), PhytoF (PF1) or vehicle alone (Ctrl) in 1% FBS media for 48 h. Cell density was later determined with crystal violet stain. All treatment values were normalized against the Ctrl group. The following symbols denote a statistical significance when compared to the control group: *, *p* < 0.05; # *p* < 0.01.

### PP1 prevents FBS-stimulated growth of MCF-7 cells

In the next experiment, the observed cytotoxic effects of PP1, PP5 and PP6 were further evaluated in the regulation of cellular proliferation of MCF-7 and MDA-MB-231. For each experiment, three 96-well plates were seeded with 10^3^ cells. 24 h after seeding, culture media was replaced with 0.1% faf-BSA for 24 h. One plate was then selected for cell density assay (0-h time point) and the others were used for exposure to PhytoPs in the presence of 10% FBS for 48 and 72 h. Control group received vehicle alone. After the end of the incubation, plates were processed for cell density assay. In order to determine cellular proliferation, 48 h and 72 h values were normalized against the 0-h values. As shown in **Figure 3**, PP5 did not prevent FBS-mediated growth of MCF-7 but only significantly affected the proliferation of MDA-MB-231 at 48 h. On the other hand, PP6 significantly inhibited growth of MCF-7 and MDA-MB-231 cells at 48 and 72 h, respectively. Our data also show that only PP1 could prevent FBS-stimulated proliferation of MCF-7 cells at both time points (Figure 3A).

**Figure 3:**
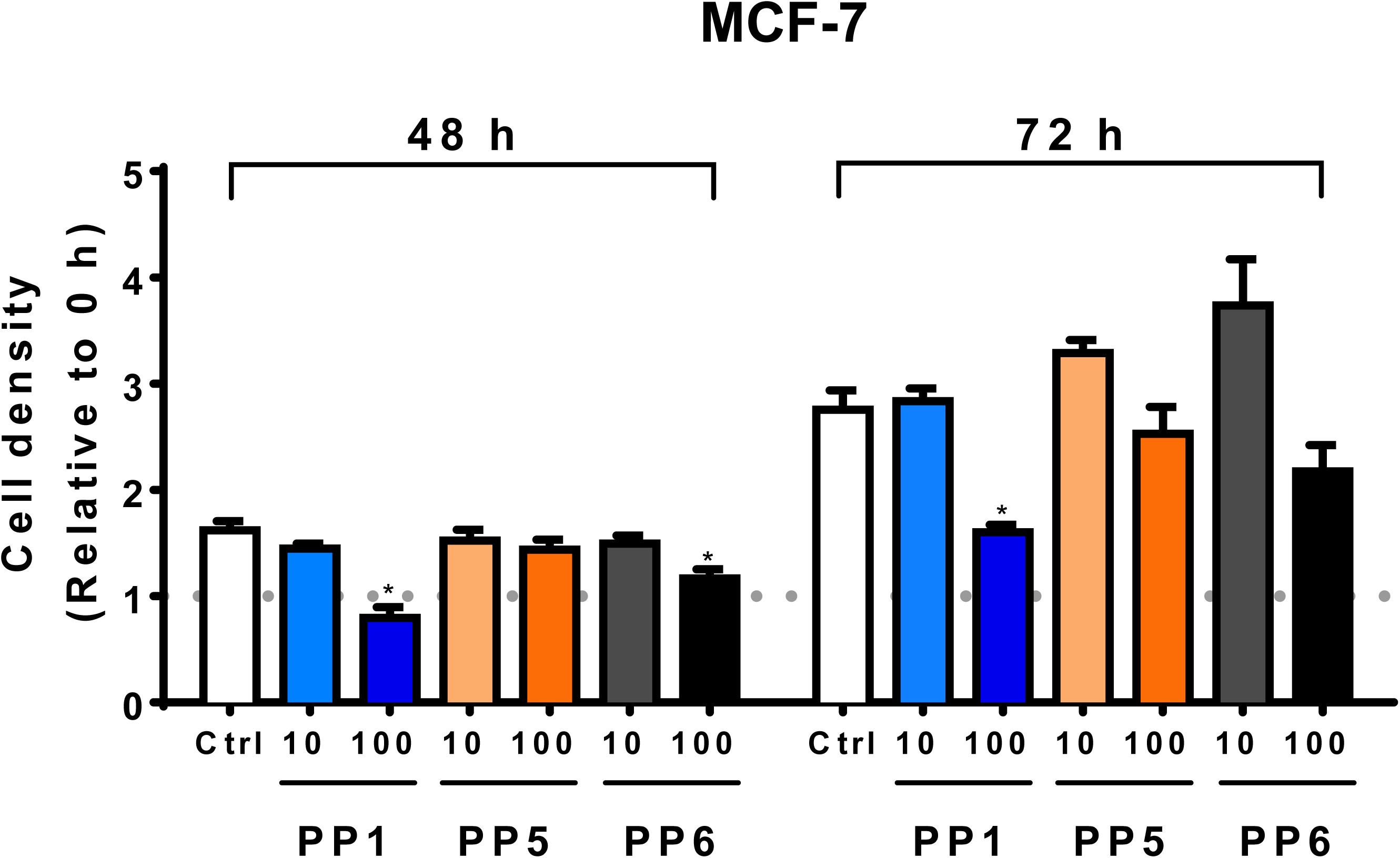
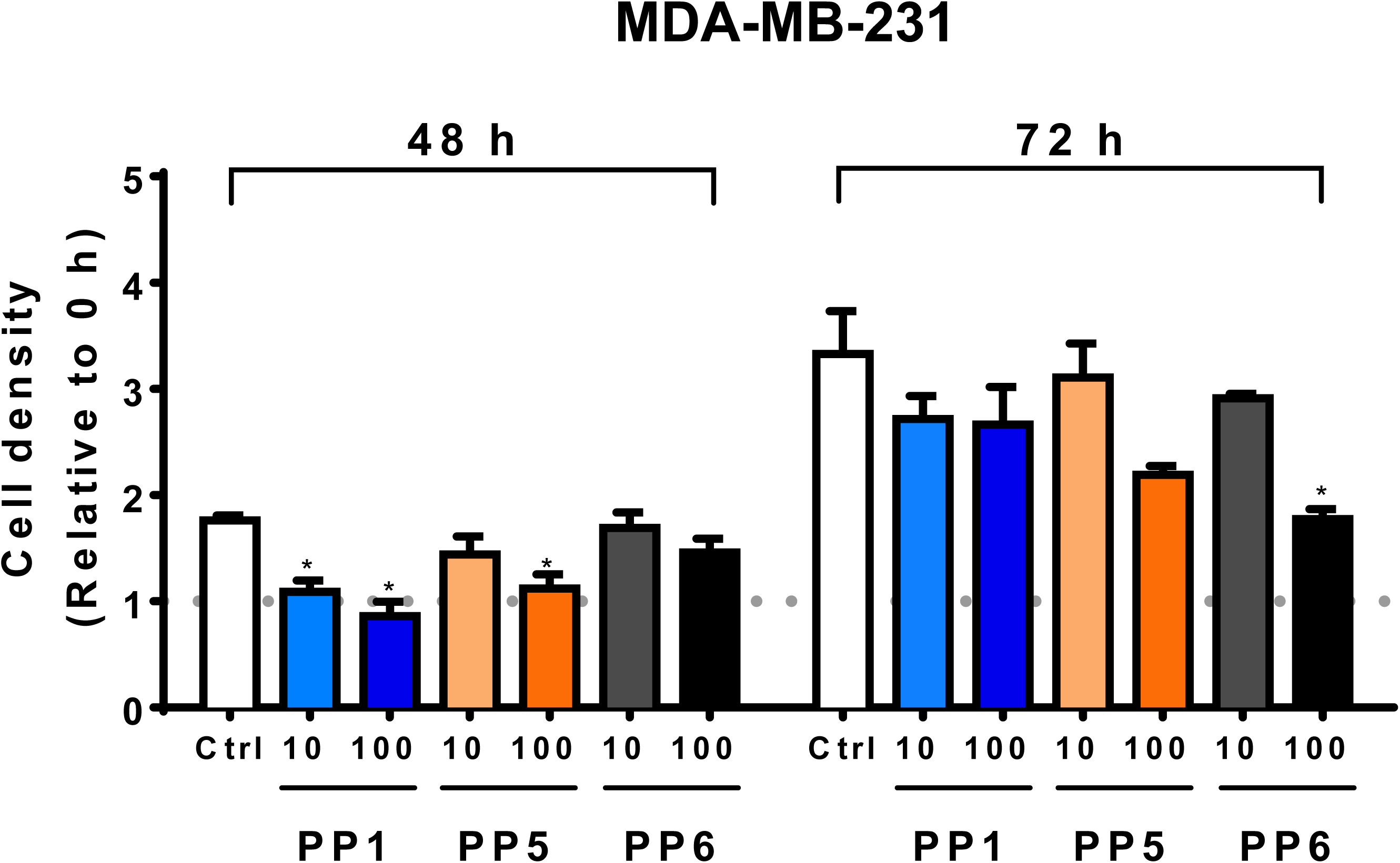
PP1 perturbs the proliferation of MCF-7 (A) and MDA-MB-231 (B) cells. 10^3^ cells per well were seeded in three 96-well plates in complete media for each experiment. 24 h later the media was replaced with 0.1% faf-BSA media and incubated for 24 h. One plate was used to determine cell density at the 0 h time point and the others were treated with vehicle alone (Ctrl), PP1, PP5 or PP6 in 10% FBS media for 48 or 72 h. At the corresponding time points, plates were subjected to a cell density assay. All treatment values were normalized against the 0-h group. The following symbol denotes a statistical significance when compared to the control group: *, *p* < 0.05.

### PP5 and PP6 block MCF-7 cells in G_0_/G_1_ and increase the number of MDA-MB-231 cells in sub G0/G1

We also examined whether PP1, PP5, and PP6 could play a role in the regulation of cell cycle progression. It was observed that PP1 could slightly increase the percentage of cells in S phase compared to control groups for both cell lines (**Figure 4**). On the other hand, incubation with PP5 and PP6 was associated with an increase in the percentage of MCF-7 cells in G0/G1 while a small percentage of MDA-MB-231 cells in sub-G0/G1 was detected. These data suggest that PP5 and PP6 could induce apoptosis of the breast cancer cells used in the present study.

**Figure 4:**
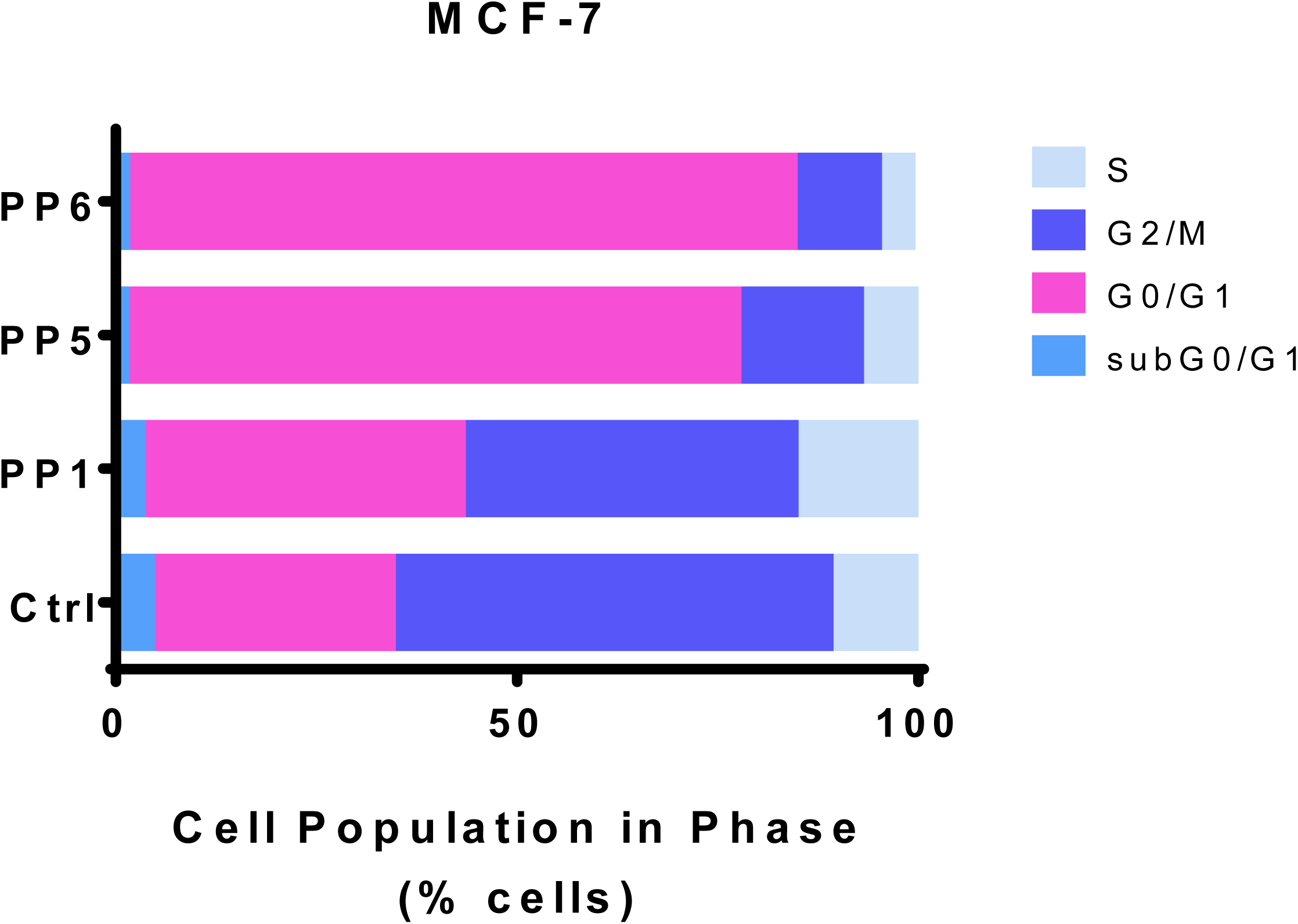
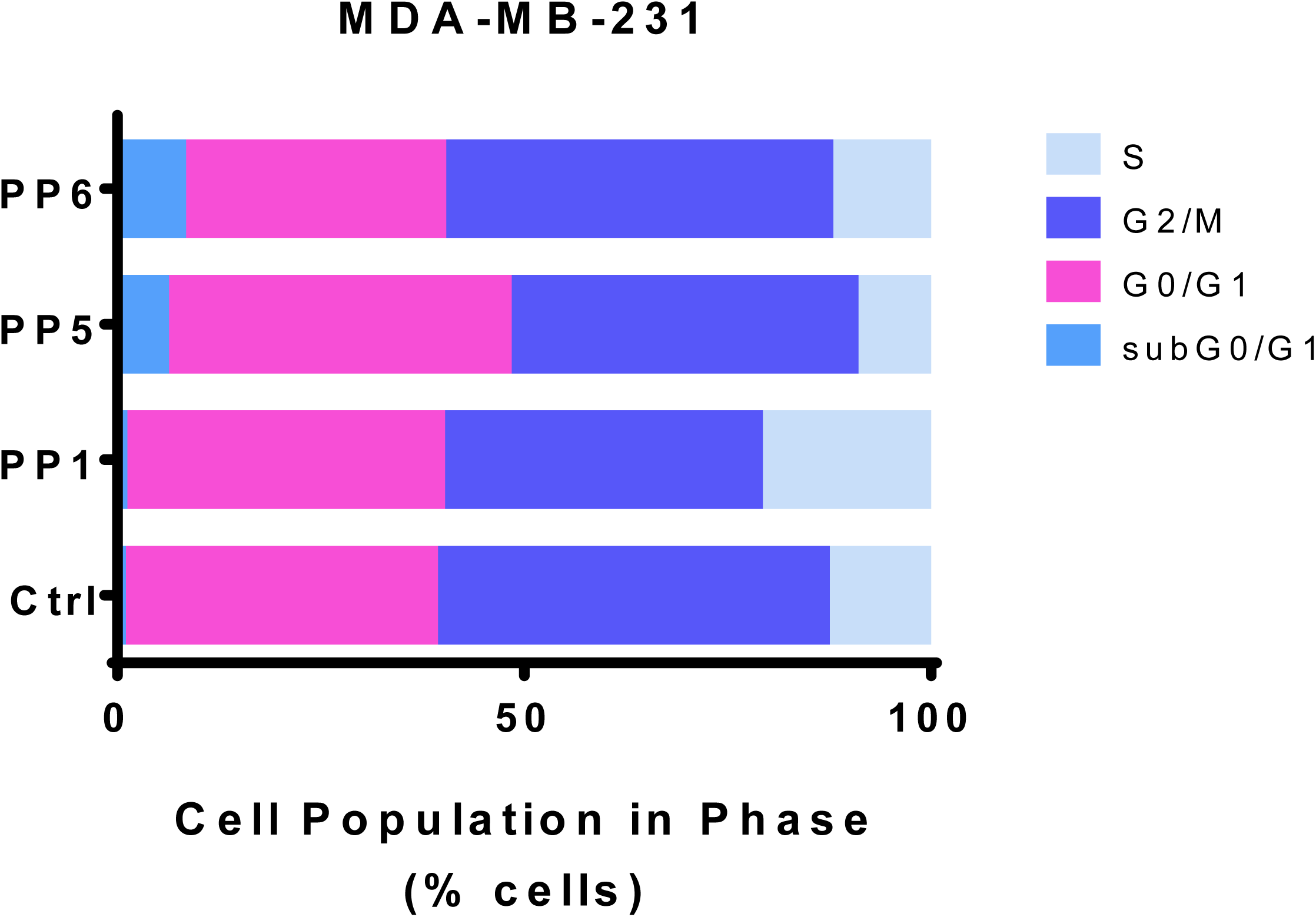
PP5 and PP6 block MCF-7 cells in G0/G1 and increase the proportion of MDA-MB-231 cells in sub G0/G1. MCF-7 (**A**) and MDA-MB-231 (**B**) (2.5×10^5^ cells/well) were seeded in 6-well-plates in complete media. 24 h later the media was replaced with 1% FBS with 100 μM PhytoPs or vehicle alone (Ctrl) and incubated for 48 h. Cells were then subjected to cell cycle analysis as described the Material and Methods section.

### PP1, in combination with a sub-cytotoxic dose of doxorubicin, reduces cellular survival

To assess the existence of a possible interaction effect between PhytoPs and a chemotherapeutic agent, MCF-7 and MDA-MB-231 were incubated with PP1, PP5, or PP6 and a sub-toxic concentration of doxorubicin, together (**Figure 5A-B**) or sequentially (**Figure 5C-D**). Our data show that, in MCF-7 cells, co-exposure of PP1 with doxorubicin could reduce cellular survival. However, this effect was not observed in MDA-MB-231 cells. Also, sequential exposure to both compounds did not improve the cytotoxicity of doxorubicin in these cell lines. These data suggest that PP1 and doxorubicin may have synergistic effects on MCF-7 proliferation.

**Figure 5:**
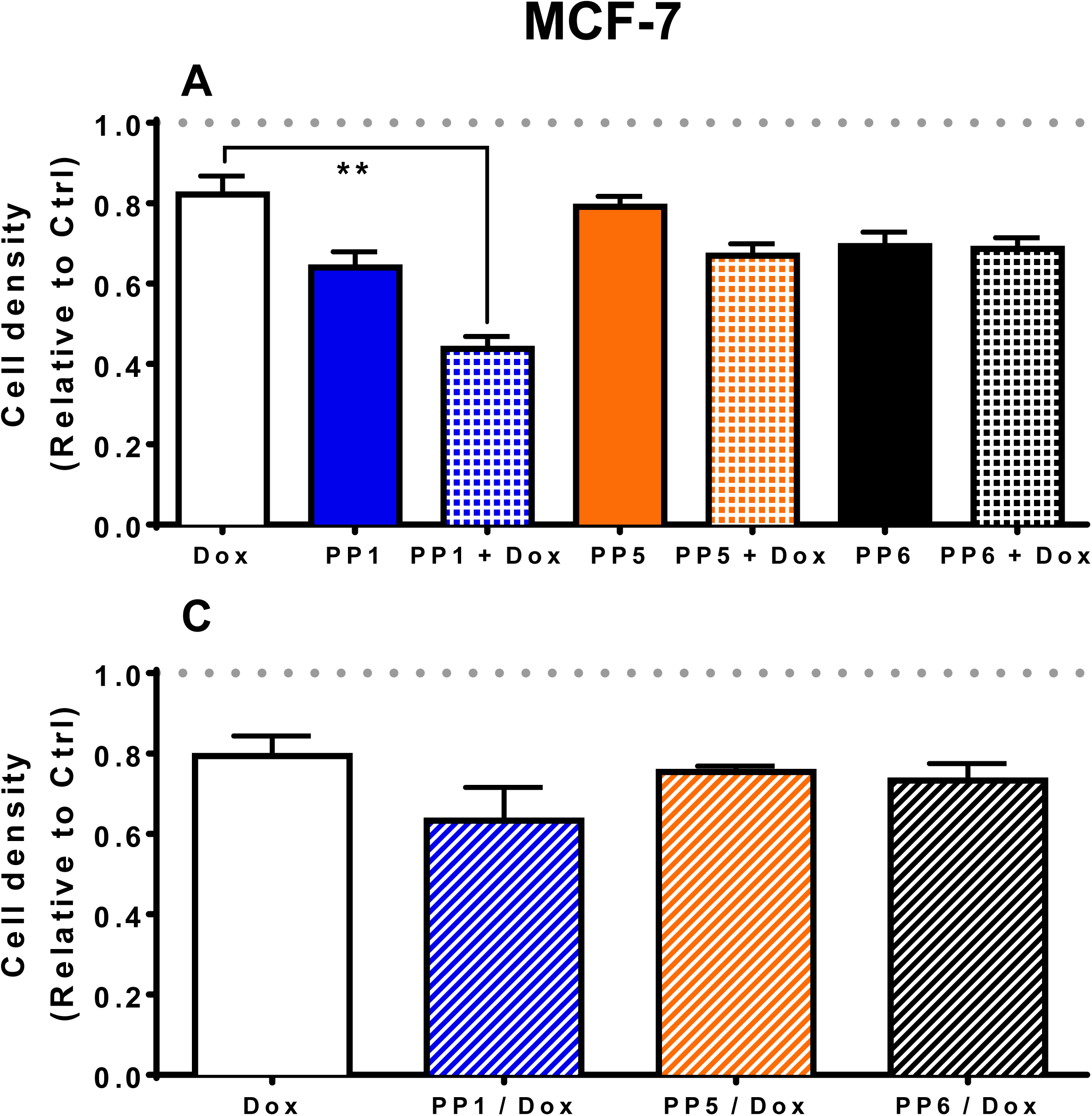
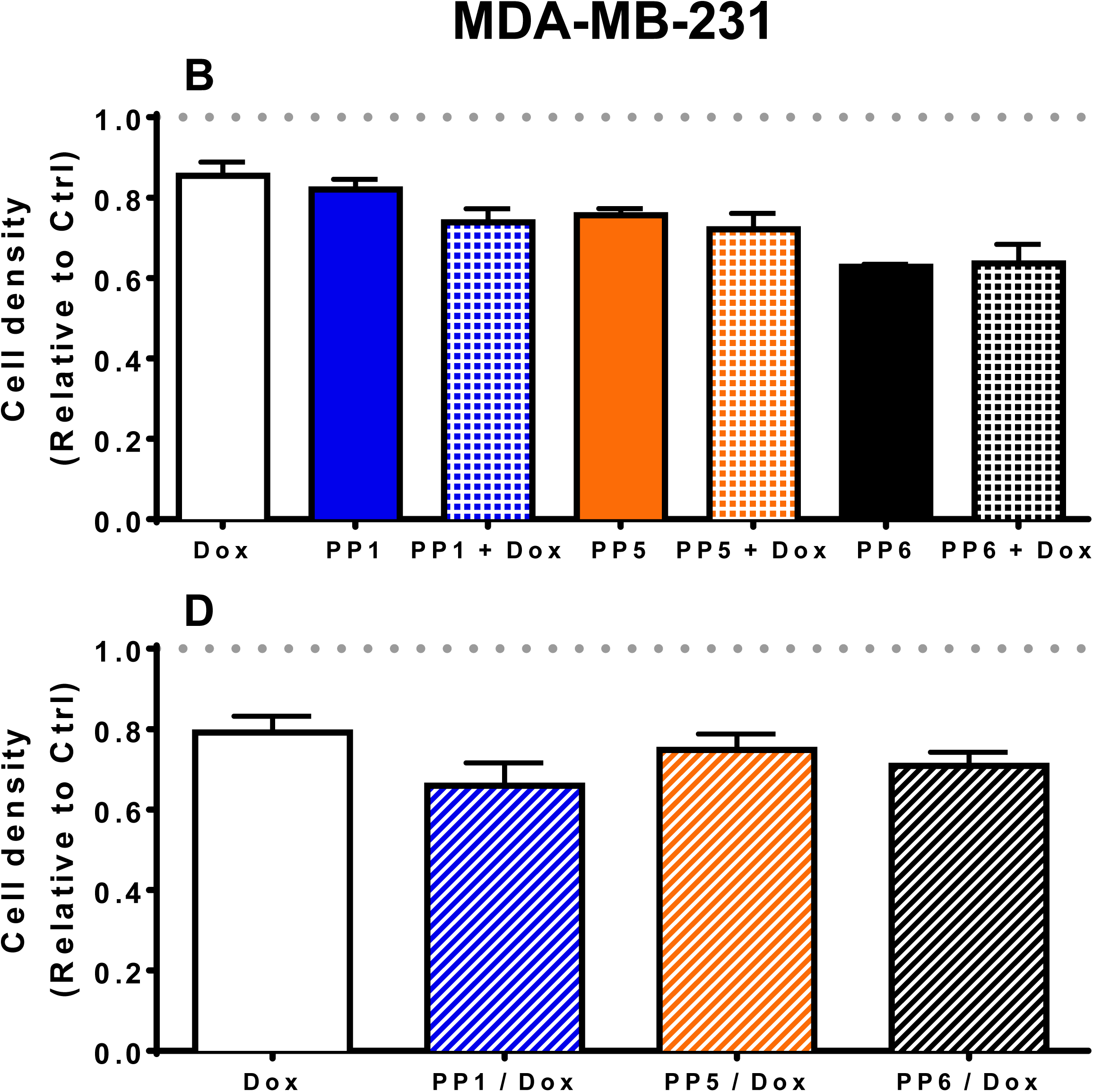
PP1 in combination with a sub-cytotoxic dose of doxorubicin reduces cell survival. MCF-7 (**A and C**) and MDA-MB-231 (**B and D**) cells were seeded in 96-well plates in complete media. 24 h later the media was replaced with 1% FBS containing 100 μM PhytoPs (PP1, PP5 or PP6) and a sub-cytotoxic dose of doxorubicin (20 nM, Dox) for 48 h. At the end of the incubation, cells were submitted to a cell density assay. (**A and B**) Results of combined exposure for 48 h are shown. (**C and D**) These panels show results after pre-incubation with PhytoPs for 48 h and media replacement with media containing 20 nM doxorubicin for an additional 48 h. Cell density was later determined with crystal violet stain. Control group (Ctrl) was incubated with vehicle alone. All treatment values were normalized against Ctrl values. The following symbol denotes a statistical significance when compared to the control group: *, *p* < 0.05.

### PF1 inhibits FBS-stimulated wound healing of MDA-MB-231 cells

A preliminary test with PP1/2/3/4/6/7 and PF1/PP2 was established to determine the potential compounds with anti-migratory effect (not shown). PP5 was excluded from this initial test due to its effect on cellular survival (**Figure 2**). Among these compounds, only PF1 displayed an effect on cellular migration, and PF1 was compared to PP2 from the same family of compounds. It was observed that the presence of 50 µM PF1 significantly retarded the migration of MDA-MB-231 cells, while PP2 had no significant effect (Figure 6).

**Figure 6:**
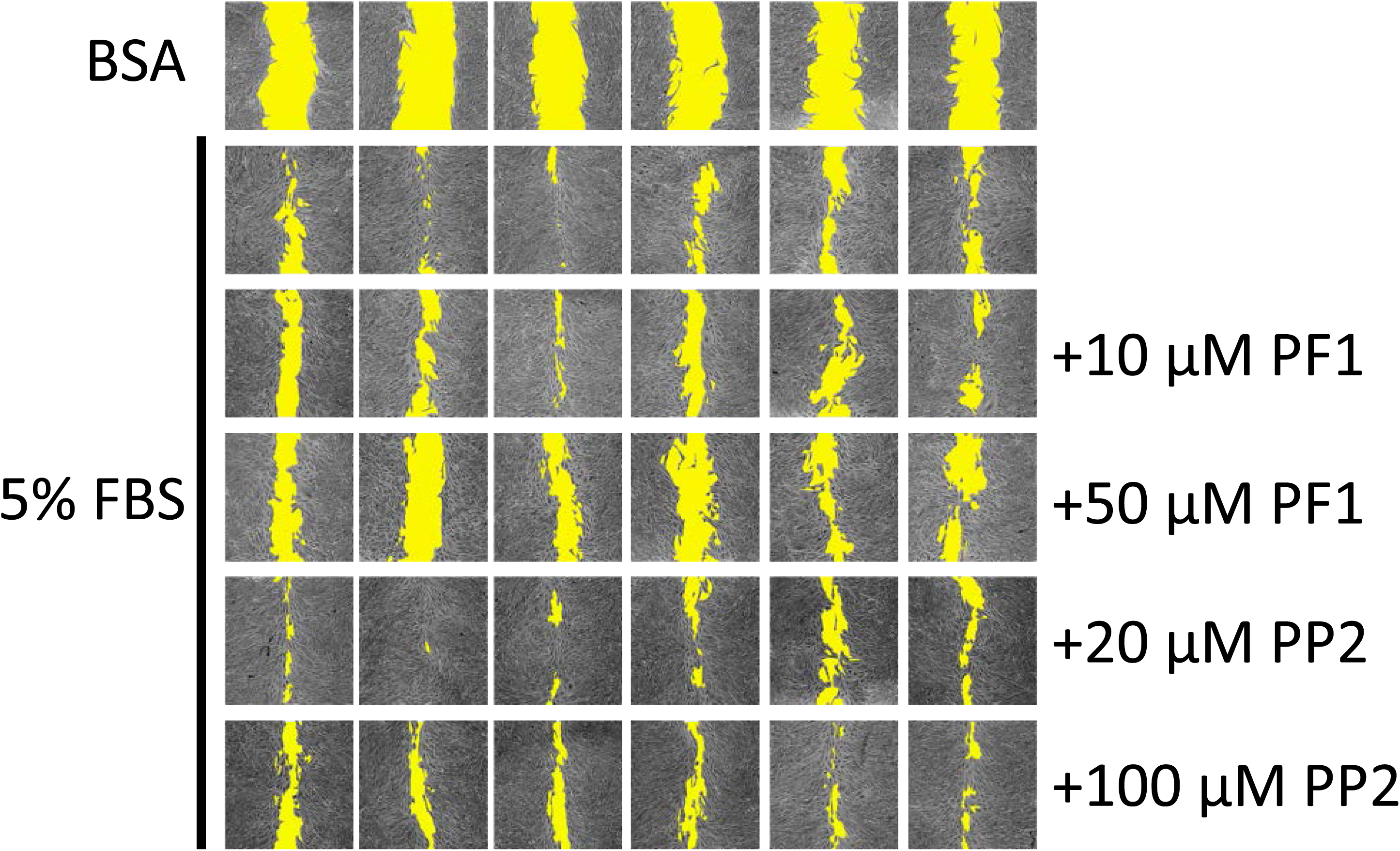
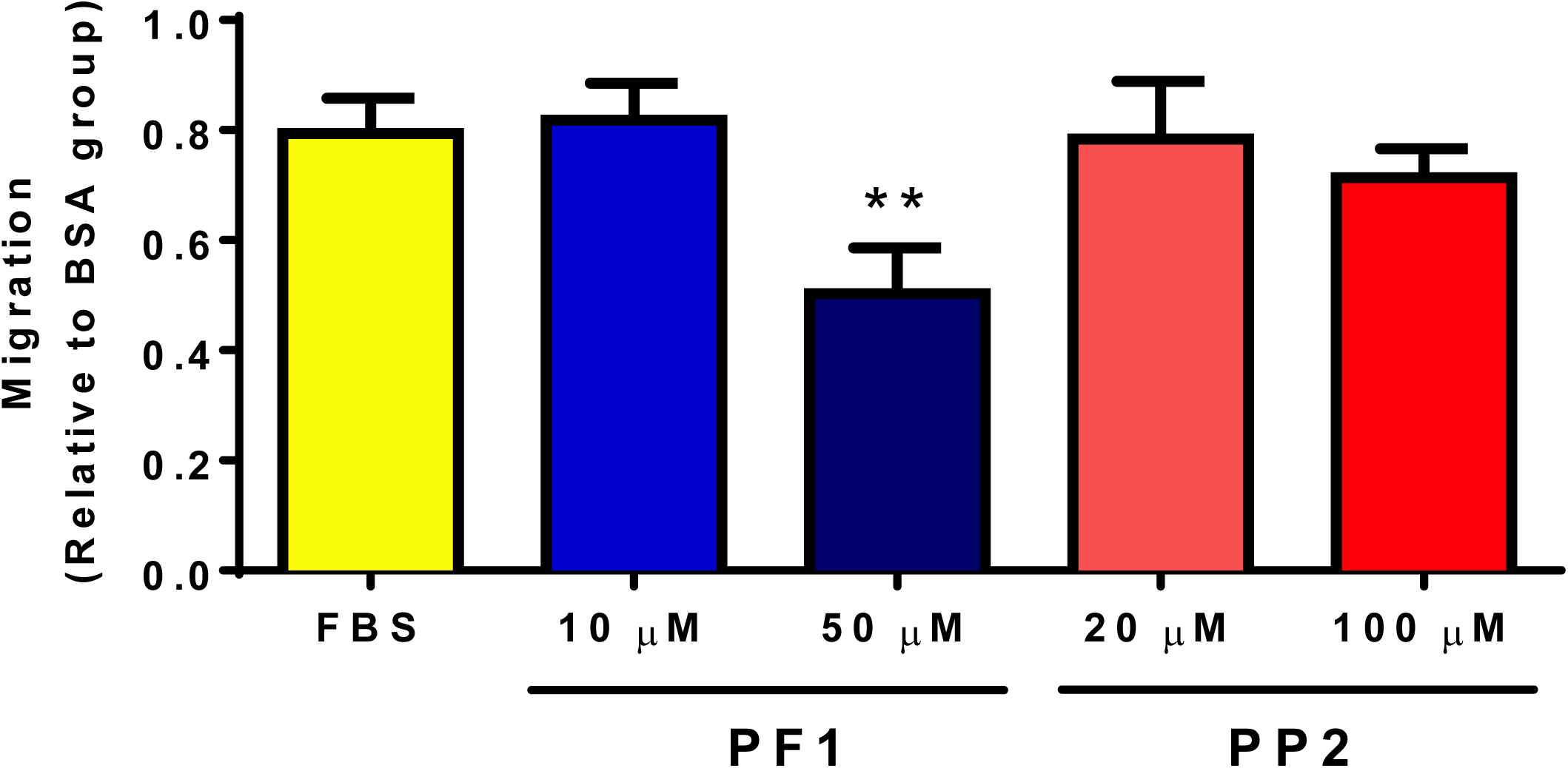
PF1 inhibits FBS-stimulated wound healing of MDA-MB-231 cells. MDA-MB-231 cells (12.5×10^3^ cells/well) were seeded in 96-well plates in complete media. Once they reached 100% confluence, complete media was replaced with media containing 0.1% faf-BSA for 24 h. A wound was then applied and media was replaced with media containing or not PF1 or PP2 30 min before adding FBS to a final concentration of 5% (v/v). Panel **A** shows representative images of wound-healing assays after 24 h. Cell-free areas are highlighted in yellow. Panel **B** shows comparison of cellular migration. The following symbol denotes a statistical significance when compared to the control group: **, *p* < 0.01.

### PF1 promotes the adhesion of MDA-MB-231 cells

To determine if PF1 could regulate cellular adhesion, PF1-pre-treated MDA-MB-231 cells were seeded on ECM-coated or uncoated plates and cellular adhesion was measured by crystal violet staining. Cellular adhesion was allowed for 1 h. As expected, barely any cellular adhesion was observed in uncoated plates. However, PF1 promoted cell adhesion of MDA-MB-231 cells on coated plates in a dose-dependent manner (**Figure 7**).

**Figure 7:**
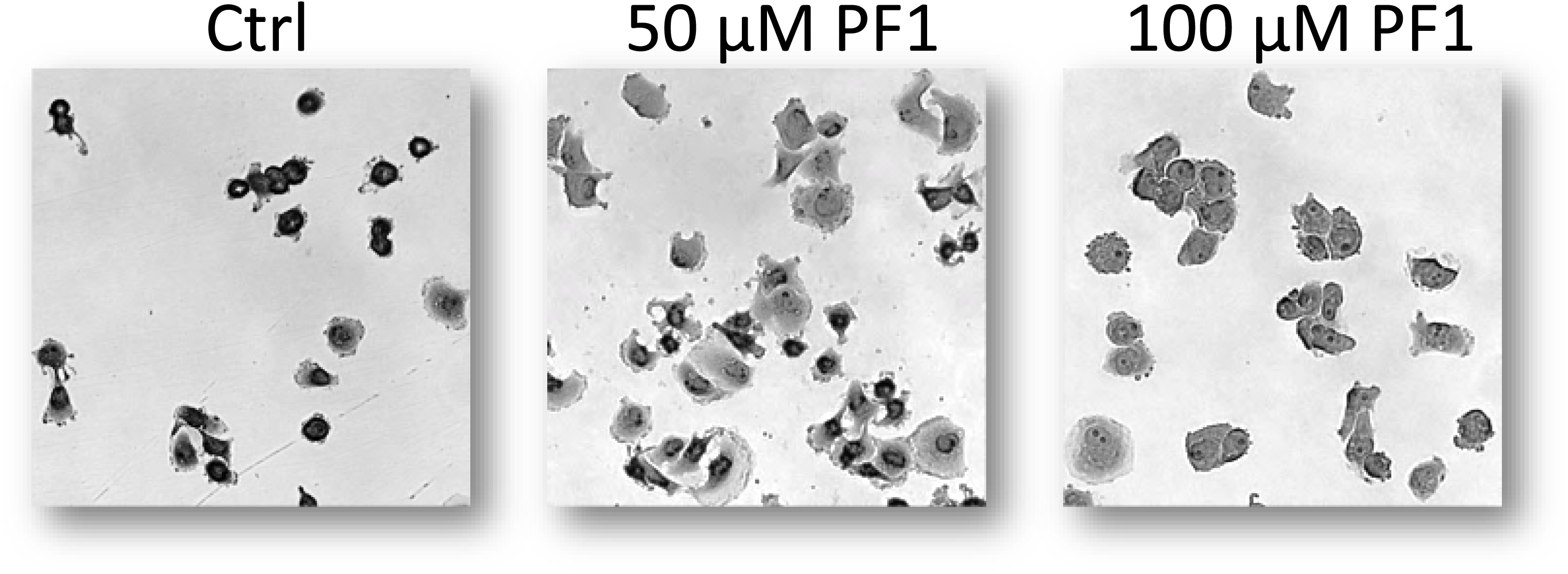
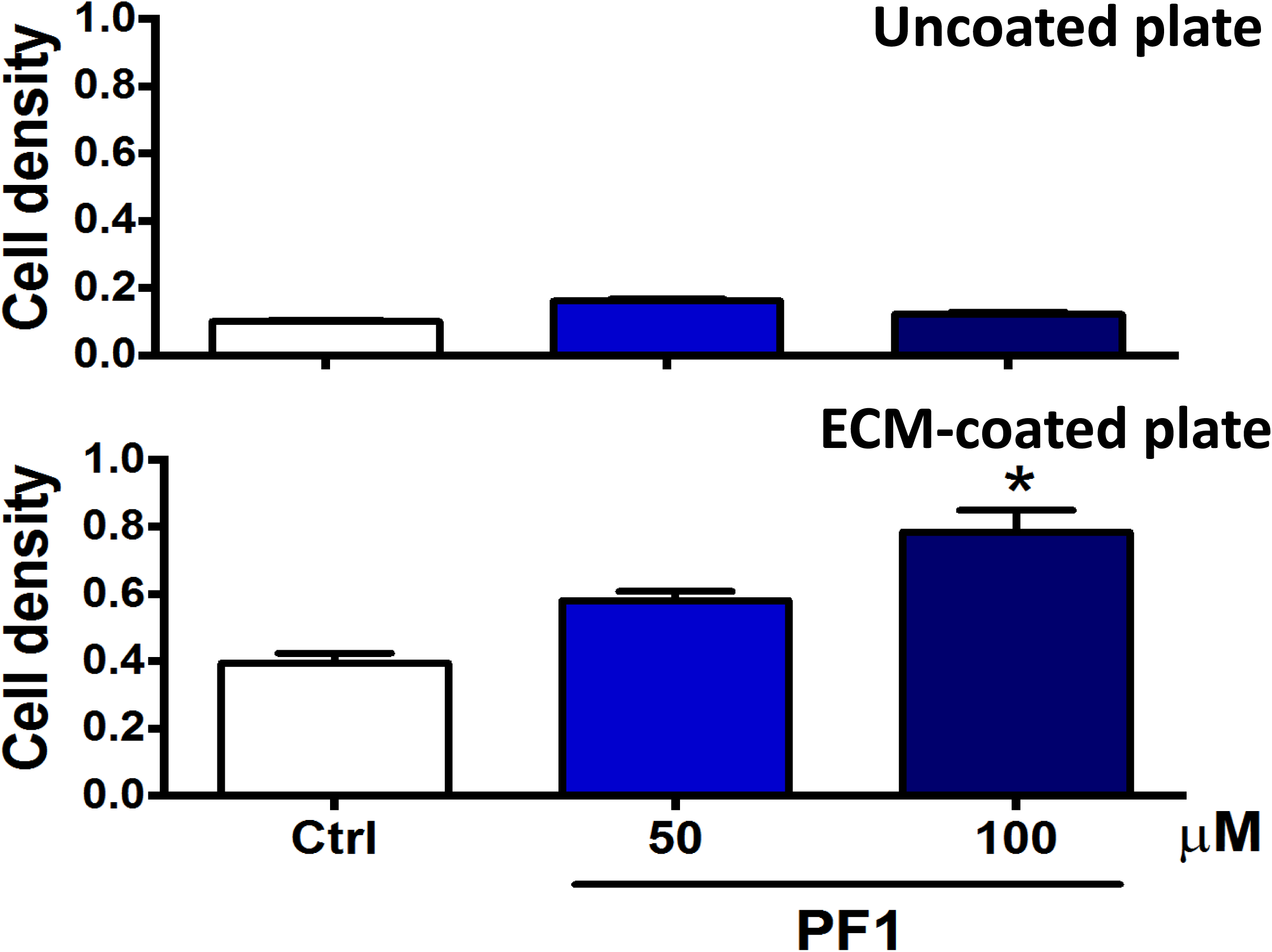
PF1 promotes the adhesion of MDA-MB-231 cells. MDA-MB-231 cells (4 × 10^6^ cells/well) were seeded in a 6-well plate. The following day, media was replaced with media containing 1% FBS and supplemented with vehicle alone (Ctrl), 50, or 100 μM PF1. After 24 h of incubation, cells were detached and seeded on uncoated (plastic) or pre-ECM-coated 96-well plates. Cellular adhesion was allowed for 1 h at 37 °C, and cells were then stained with crystal violet. After drying, pictures were taken before quantification. Panel **A** shows cells seeded on ECM-coated plate. Note the increased number of well-spread cells when treated with PF1. Panel **B** shows quantification of intracellular CV. The following symbol denotes a statistical significance when compared to the control group: *, *p* < 0.05.

## Discussion

In the present work, eight non-enzymatically oxidized derivatives of ALA were tested to identify their potential anticancer activities. Among them, we have identified that 16-F_1t_-PhytoP (PP1) and *Ent*-9-L_1_-PhytoP (PP6) can regulate cellular survival and proliferation of breast cancer cell lines. Also, *Ent*-9-(*RS*)-12-*epi*-ST-Δ^10^-13-PhytoF (PF1) displays anti-migratory activity with MDA-MB-231 cells. These findings are important and suggest that these compounds may be usable for the treatment of breast cancer, and they may be used for the treatment of primary tumor as well as metastasis development. However, further evaluation will be required.

While AA is the source of IsoPs in animals, ALA is the source of PhytoPs and PhytoFs in plants since AA availability in plants is limited. Interestingly, in patients fed a diet enriched in flaxseed oil, increased plasma levels of PhytoPs have been observed [15]. While some of the PhytoPs may be directly derived from the diet [12], data show that ALA may be transformed into PhytoPs in the human body [15]. These findings suggest that PhytoP formation may be relevant *in vivo* and may have specific effects in human diseases. Studies have shown that, in breast cancer, ALA limits tumor progression by inhibiting cancer cell proliferation [16, 17] and tumor progression [18, 19]. These effects may be mediated by other n-3 fatty acids (EPA and DHA) but independent roles for ALA have been proposed [20, 21]. Derivatives of ALA such as PhytoPs and PhytoFs may therefore play a specific role in the inhibition of breast cancer progression mediated by ALA.

Structurally, PhytoPs, PhytoFs, and IsoPs are related. Therefore, their biological effects may also be overlapping. Studies have shown that 15-F_2t_-IsoP, 15-E_2t_-IsoP and 15-A_2t_-IsoP inhibit VEGF-induced migration, tube formation by ECs, and cardiac angiogenesis *in vitro*, as well as VEGF-induced angiogenesis *in vivo* via activation of the thromboxane A2 receptor (TBXA2R) [13]. Notably, an increased expression of TBXA2R was observed in tumors of human breast cancer [25], and activation of this receptor promoted the proliferation of lung carcinoma cells [26] and survival of triple negative breast cancer cells [27]. Therefore, it is reasonable to propose that the observed effects of PhytoPs and PhytoFs on cellular survival and proliferation in the present work could be related to their ability to regulate TBXA2R function in breast cancer cells. In addition, TBXA2R has also been involved in the migration and invasion of breast cancer cells [27, 28]. Taken together, these data suggest that ALA and ARA oxidative derivatives may have distinct and possibly opposite effects on cancer cell properties that may be, at least in part, mediated by TBXA2R.

Other pathways regulating cancer cell migration may also be involved in the effect mediated by PF1. As shown in this work, PF1 could diminish MDA-MB-231 response to serum-stimulated migration in a wound-healing assay (**Figure 6**). This effect could be explained, at least in part, by the increased adherence of cells exposed to PF1 (**Figure 7**). An inverse relationship between cellular adhesion and migration has previously been described for pancreatic cancer cells where down-regulation of p8 promotes cellular adherence and decreases cellular migration via regulation of cdc42 [29]. This inverse relationship has also been described under hypoxic conditions for L929 fibroblasts, which increase the number of focal adhesion contacts per cell and cellular surface β1-integrin levels [30]. Although no specific study has been performed on PhytoPs or PhytoFs, IsoPs have been shown to increase adhesiveness of neutrophils [31] and platelets [32]. Further studies are required to establish the molecular pathways involved in PF1 regulation of cellular migration.

Alternatively, one type of PhytoPs has also been identified as an activator of nuclear factor-erythroid 2-related factor 2 (Nrf2), which is a transcription factor that subsequently triggers a cellular oxidative stress response [33]. In that regard, other naturally occurring, plant-derived phytochemicals have been identified as activators of Nrf2. It was proposed that the chemotherapeutic activities of these compounds may be mediated by Nrf2. Nrf2 may allow the activation of phase II detoxification enzymes, antioxidants, and transporters [34].

## Conclusions

Given the fact that the human body can produce PhytoPs and PhytoFs, consumption of a diet enriched in ALA may affect cancer progression and/or response to chemotherapeutic treatment, at least in the case of breast cancer. In addition, it may also attenuate inflammation, as previously shown. Taken together, these findings are important and suggest that these compounds may be usable for the treatment of breast cancer, and they may be used for the treatment of primary tumor as well as metastasis development. However, further evaluation will be required.

## Declarations

### Availability of data and materials

All data generated or analyzed during this study are included in this published article.

### Competing interests

The authors declare that they have no competing interests.

### Funding

JLGP was supported by Le Studium (Région Centre-Val de Loire, France). PGF was supported by grants from INCa PLBio (2018-145), the Lipids ARD2020-Biodrug project (Région Centre-Val de Loire, France), La Ligue contre le Cancer (Indre et Loire, Loir et Cher, and Vienne), by an Academic Research Grant from the Région Centre-Val de Loire (France) and by the Canceropole Grand-Ouest (Mature project). The authors thank Valérie Bultel-Poncé and Alexandre Guy from IBMM for their participation in the synthesis of PhytoPs and PhytoFs.

### Authors’ contributions

JLGP and PGF designed research. JLGP, CBH, CO, JMG, TD, and PGF performed research; and JLGP and PGF wrote the paper. All authors have read and approved the manuscript.

## Notes

#### Summary of Updates

Abstract: corrections Figure 1: revised Typo corrections

